# DRUG-seq Provides Unbiased Biological Activity Readouts for Drug Discovery

**DOI:** 10.1101/2021.06.07.447456

**Authors:** Jingyao Li, Daniel J. Ho, Martin Henault, Chian Yang, Marilisa Neri, Robin Ge, Steffen Renner, Leandra Mansur, Alicia Lindeman, Tayfun Tumkaya, Carsten Russ, Marc Hild, Caroline Gubser Keller, Jeremy L. Jenkins, Kathleen A. Worringer, Frederic D. Sigoillot, Robert J. Ihry

**Affiliations:** Neuroscience, Novartis Institutes for BioMedical Research, Cambridge, MA; Chemical and Biological Therapeutics, Novartis Institutes for BioMedical Research, Cambridge, MA

## Abstract

Unbiased transcriptomic RNA-seq data has provided deep insights about biological processes. However, its impact in drug discovery has been narrow given high costs and low throughput. Proof-of-concept studies with Digital RNA with pertUrbation of Genes (DRUG)-seq demonstrated the potential to address this gap. We extended the DRUG-seq platform by subjecting it to rigorous testing and by adding an open-source analysis pipeline. The results demonstrate high reproducibility and ability to resolve the mechanism(s) of action for a diverse set of compounds. Overall, the protocol and open-source analysis pipeline are a step towards industrializing RNA-seq for high complexity transcriptomics studies performed at a saturating scale.

## Introduction

In the pharmaceutical industry, it is standard to test thousands of compounds in high-throughput screens to identify regulators of a target or a biological process (Swinney and Anthony, 2011; Volochnyuk et al., 2019). This massive scale is made possible by focusing on a single readout. However, biological systems are inherently complex, and there is a need for scalable screening methods that can capture the total biological activity of small molecule libraries. Whole-transcriptome analysis, by RNA-seq, offers a high-dimensional readout but is cost prohibitive, and is typically performed on a small number of samples (Cleary et al., 2017; Verbist and Horchreiter, 2015).

To reduce costs, targeted RNA-seq approaches such as L1000 or RASL-seq have been deployed successfully as large-scale transcriptomic profiling methods (Li et al., 2012; Subramanian et al., 2017). Targeted approaches such as these can be tailored to any gene set of interest, but it takes time to optimize gene sets for any given disease or cell model (Kong et al., 2021), and this approach may miss unexpected and potentially important transcriptomic signatures. Single cell RNA-seq can detect a few thousand genes per cell in an unbiased way. Recently the sci-Plex method identified cell-specific transcriptional responses to hundreds of compounds by labeling the cells in each treated well with a single-stranded DNA barcode that binds to nuclei prior to single cell (sc)RNA-seq sample processing, which enables sample multiplexing (Srivatsan et al., 2020). It will be promising to apply methods like sci-Plex to study the single cell effects of treatment in complex tissue models, such as brain organoids (Lancaster and Knoblich, 2014). As cell diversity increases, 1000s of cells must be profiled to detect responses in lower abundance or rare cell types. For more homogeneous cell culture models, scRNA-seq plus perturbation methods may offer less of an advantage over bulk RNA-seq methods and cellular resolution comes at the expense of throughput of perturbations tested and the number of genes detected.

DRUG-seq is a low-cost, high throughput bulk RNA-seq method that uses a direct in-well lysis of cells in 384-well plates and is ideal for studying the transcriptomic effect of many compound treatments in parallel (Ye et al., 2018). Multiple groups have been working to develop unbiased whole transcriptome-wide methods that are lower cost (Bush et al., 2017; Sholder et al., 2020; Yeakley et al., 2017). The pace of development in this field makes it difficult to compare data or perform exhaustive benchmarking studies to compare performance of the methods. As a key step toward this, we created an experimental design to thoroughly test the DRUG-seq platform and made the methods, data, and code available for independent verification and reproduction of the results. By exhaustively testing reproducibility across batches and plates, we demonstrate that DRUG-seq provides the granularity to bin compounds by mechanism of action (MoA) and meets the performance standards required of an industry-scale RNA-seq platform.

## Results

### Summary of Protocol and Experimental Design

DRUG-seq pairs high-throughput cell culture with miniaturized RNA-seq (Figure 1A). Cells are first cultured in 384-well plates and then treated with compounds for a desired period of time. After the treatment, the cells are directly lysed in each well, without RNA purification. The RNA in each well is used as a template for reverse transcription (RT), where the DRUG-seq RT primers incorporate both the well barcodes and unique molecular identifiers (UMI) (Figure 1B). After RT, the samples are pooled and used as a template for second strand cDNA synthesis and subsequent library construction. During library construction, Tn5-mediated cDNA tagmentation is performed which enzymatically fragments and adds Illumina adaptors to each insert. Next plate-level barcodes are incorporated to track multiple plates simultaneously. Finally, for optimal sequencing, libraries are size-selected, and quality checked by DNA fragment analysis. After quantification, libraries for each plate(s) are normalized and pooled, such that we can accurately target sequencing at a read depth equivalent to 1 million reads per well on the Illumina platform. The final product is 3’ biased; full-length transcripts are not sequenced. The protocol scale can be adapted depending on the experimental need and resources available.

**Figure 1.**
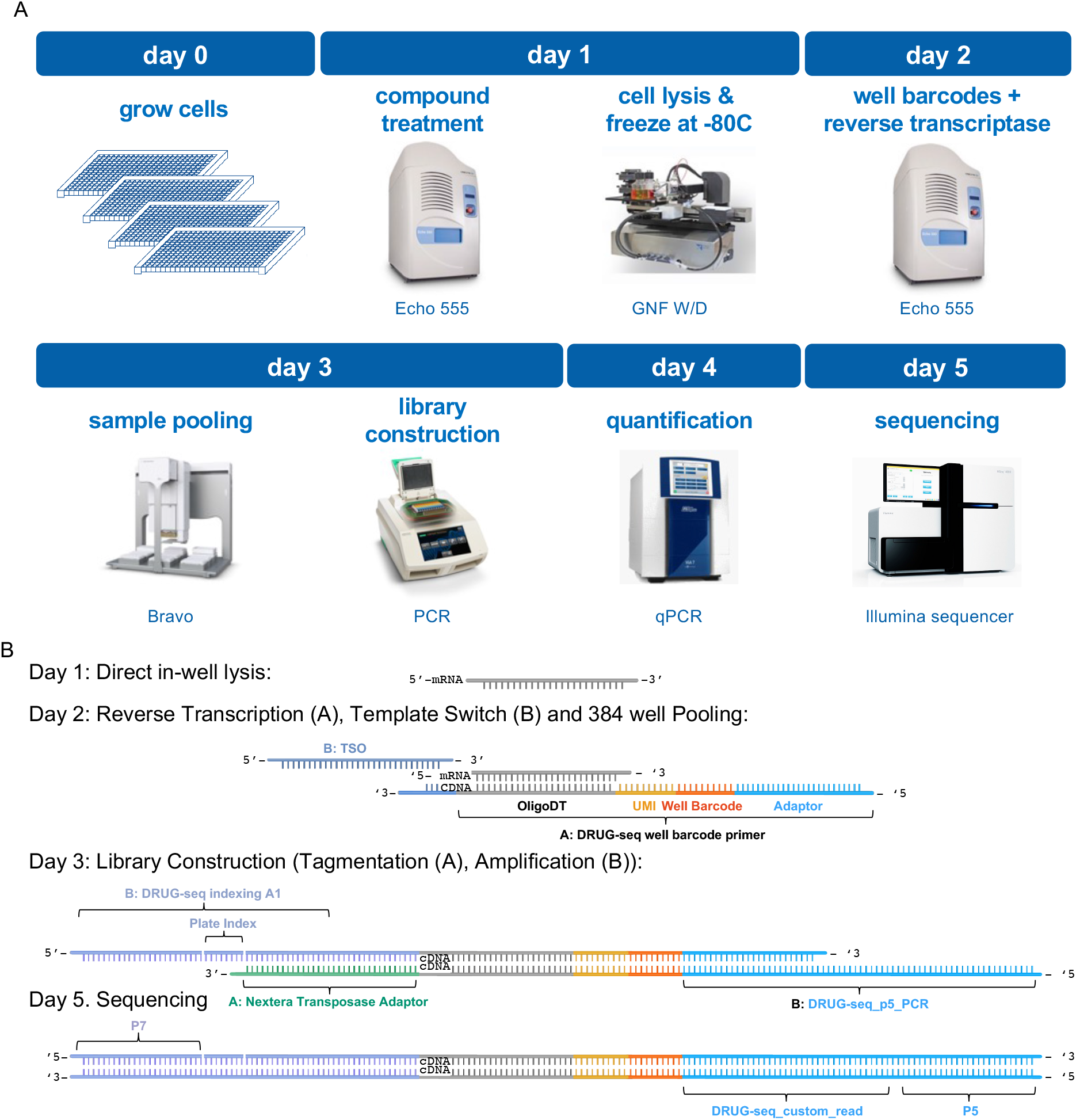
Overview of the DRUG-seq protocol. (A) Once cells are plated and treated with perturbagens the DRUG-seq protocol can be performed and sequenced in 5 days. The process and equipment depicted here is for batches of 18 or fewer plates. Scale up or down is possible depending upon infrastructure available. (B) Depiction of the DRUG-seq high throughput RNA-seq chemistry (Adapted from Ye et. al., 2018). Day 1: The scale of DRUG-seq is enabled by using a direct in-well lysis of cells without mRNA purification, which is coupled to a barcoding and sequencing strategy, on day 2, that tracks transcripts (UMI), wells (well barcode), and plates of cDNA libraries (Illumina Nextera or custom Illumina indexes) sequenced on the Illumina platform (days 3-5).

We designed an experiment to evaluate the reproducibility of the DRUG-seq protocol (Figure 2A). We generated 3 independent batches of U-2 OS cells on different days, plated in 3 replicate 384-well plates per batch for a total of 9x 384-well plates. In each plate, we treated cells with 14 compounds (Figure 2B, File S1, Table 1), each with an 8-point dose response (3.2 nM to 10 uM) with 3 replicates for each dose. This design was used to test the performance between batches of cells and plate(s).

**Table 1.** 14-compounds.

| Compound | NSN identifier | Target | Cluster<br>Ye et. al., 2018 |
| --- | --- | --- | --- |
| Triptolide | ND-09-QY33 | ERCC3i | IV |
| CHIR118637 | CA-90-VK59 | GSK3Bi |  |
| Cmp_341 | JC-68-AL90 | JAK2i | IV |
| Fedratinib | MB-03-IE36 | JAK2i | III |
| NVS-SM2 | JC-43-ZO95 | SNRPCmod | II |
| CPI-203 | PA-33-CK45 | BRD4i | III |
| Brutasol | KA-73-NB69 | NRF2i | VI |
| Homoharringtonine | QC-05-UB63 | EIF4Ei | VI |
| BTdCPU | SE-15-AV21 | EIF2AK1a | VI |
| AZD8055 | NV-67-DX31 | MTORi | VI |
| NVP-BVB808/Cmp_334 | DB-85-YA47 | PI3K | I |
| Dilazep | NA-37-JQ34 | SLC29A2i |  |
| BLU9931 | VA-76-OV33 | FGFR4 | IV |
| (S)-Crizotinib | LD-22-SA99 | ALK | IV |

**Figure 2.**
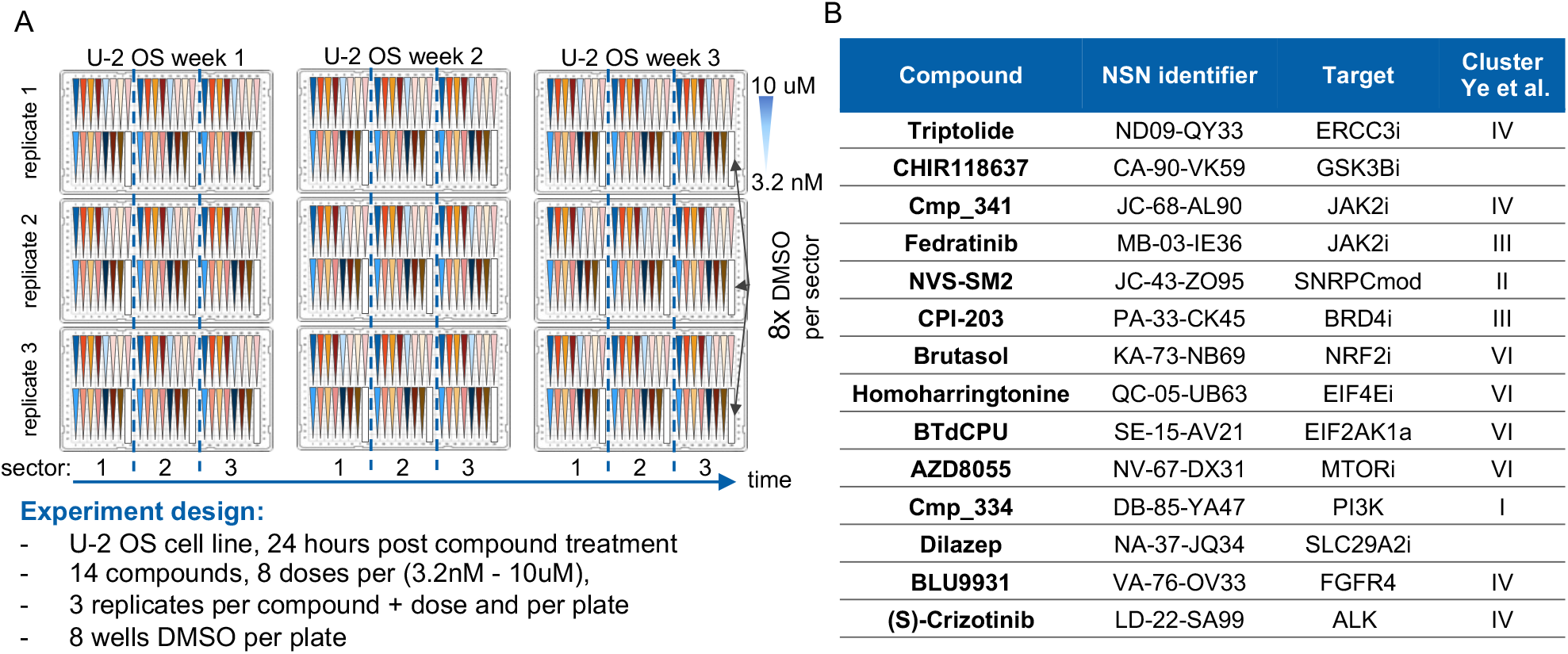
DRUG-seq reproducibility experimental design. (A) Experimental design depicted by 9 plate maps. Within each plate 14-compounds were plated in 8-doses (3.2nM-10uM) in 3-sectors for a total of 3 replicates per condition per plate (n=3/per condition). Each sector contained 8 DMSO wells for a total of 24 per plate (n=24). For each week (or batch) of cells there were 3 replicate plates. Each batch of cells were plated on independent days to reflect biological variability. (B) Table depicts compound name, identifier, target, and cluster from Ye et. al., 2018 publication. 14-compounds were selected to represent a diverse set of MoAs.

### DRUG-seq Analysis Pipeline and Activity Threshold

The data processing and primary analysis pipeline is a series of manually run steps, which allow for stepwise review and quality control. After sequencing, reads are demultiplexed using Illumina’s standard bcl2fastq2 method. The upstream DRUG-seq data processing pipeline employs highly parallelized mapping, as well as barcode and UMI counting and filtering to efficiently generate count tables (Figure 3A). Mapping of sequencing reads is performed using a custom version of the Ensembl GRCh38 reference, in which each gene was constructed using all annotated exons.

**Figure 3.**
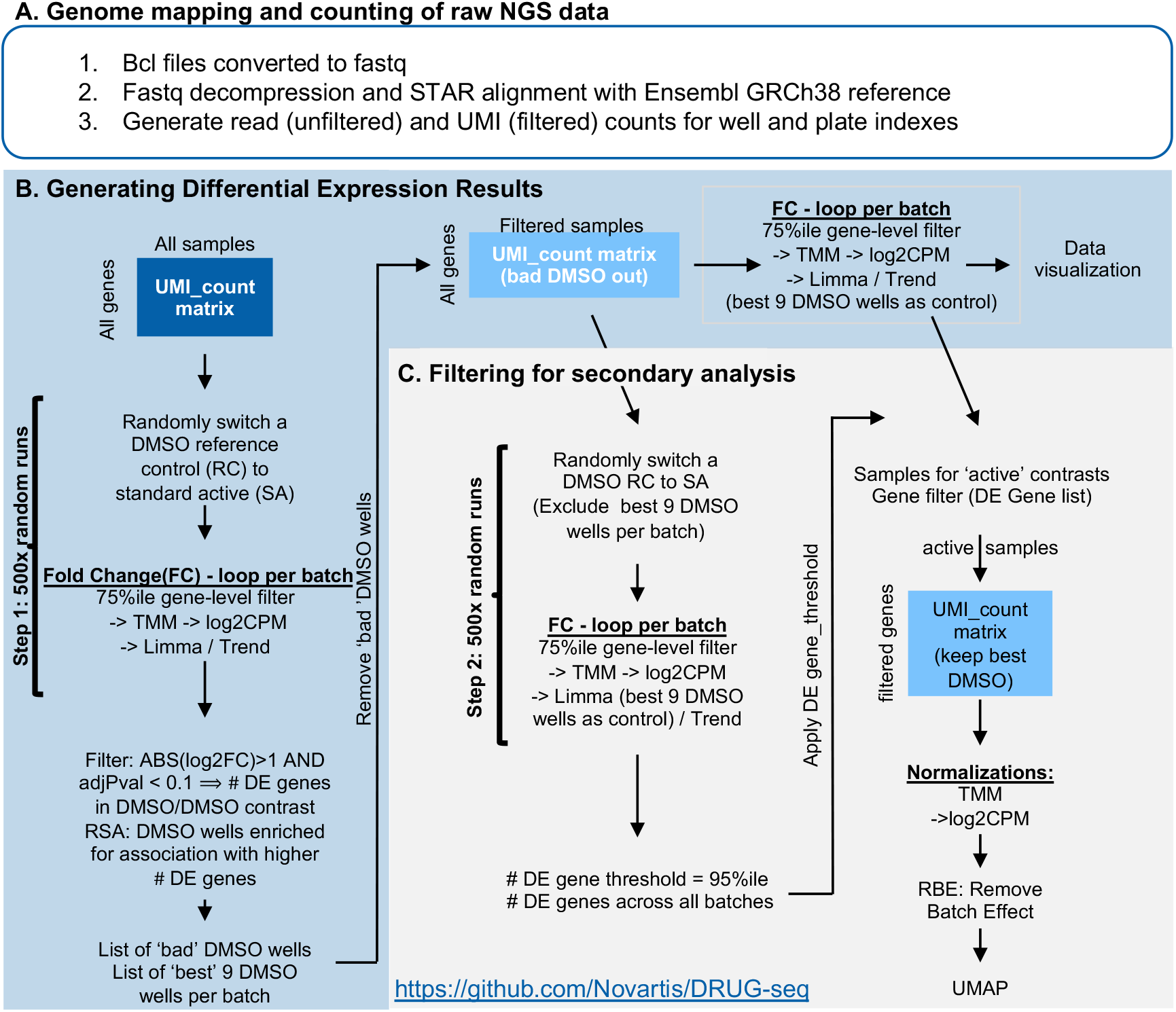
DRUG-seq analysis pipeline. The DRUG-seq analysis pipeline is comprised of 3 steps (A) Outline of the steps required to convert raw next generation sequencing data into a count matrix by converting to fastq files, aligning to the transcriptome and counting transcripts associated with well and plate barcodes. (B) Flow chart describing how differential expression results are obtained from the UMI count matrix generated in part A. In step one, the true null is calculated by performing 500 random DMSO to DMSO comparisons to generate DE results. Next, RSA analysis is used to rank DMSO wells by the number of DE genes they contributed to. Bad DMSO wells are removed and the 9 best DMSO wells are used as reference control (RC) to calculate differential expression for the compound treated wells. (C) Flow chart describing the filtering steps required for secondary analysis. Step two of the true null calculation is performed and 500 random DMSO to DMSO comparisons generate DE gene results in the absence of bad wells. The 95^th^ percentile of the 500-comparison distribution is then used as an activity threshold. The threshold is the minimal number of DE genes required to be greater than the technical noise in DMSO 95% of the time. The true null activity threshold is used to filter active samples. DE gene filtering, normalization, and removal of batch effects are applied to generate the final UMAP visualization.

The following steps of the analysis code are shared through GitHub (https://github.com/Novartis/DRUG-seq). Before generating differential expression (DE) results, an activity threshold is empirically determined based on the technical noise of the experiment and is coined the ‘true null calculation’. This takes advantage of the high number of DMSO wells that can be sampled in this low-cost RNA-seq assay. The baseline for transcriptionally active treatments is set by performing multiple permutations of randomly sampled DMSO vs. DMSO comparisons using all available wells (step one). In step one, for each batch of three plates, three random DMSO wells were selected (one per plate) and were compared to the remaining 69 DMSO wells to then calculate DE. This process was iterated 500 times (Figure 3B, 4A, File S2). It should be noted that the number of iterations, as well as number of DMSO wells chosen as optimal reference controls can be customized for each experiment. Out of the 500 iterations, we identified both DMSO wells that contributed to the fewest DE genes and DMSO wells that contributed to the most DE genes using the Redundant siRNA activity (RSA) statistic (König et al., 2007). The nine best DMSO wells per batch (three per plate) were then used as reference controls to calculate DE for the compound treated samples (Figure 3B). DE analysis is conducted using limma-trend (Ritchie et al., 2015) to quantitate gene level changes in the transcriptome, which can serve as the input for additional analyses.

In step two, DMSO wells that potentially inflate the number of DEGs were removed (‘bad’ wells marked red, Figure 4A). The true null was recalculated by comparing three random DMSO wells versus the nine best DMSO wells with 500 iterations, where we selected the three random wells from the remaining DMSO wells (Figure 3C, 4A, File S3). When comparing the results from step one and two, the frequency of DMSO to DMSO comparisons yielding more than 100 DE genes was reduced with reciprocal gains in the frequency of DMSO comparisons with 20 or fewer DE genes (Figure 4B). We typically define transcriptionally active compounds as those with more DE genes than the 95^th^ percentile of the DMSO to DMSO true null distribution (Figure 3C, 4C). This indicates the treatment is more active than DMSO 95% of the time. Results obtained with all DMSO wells (Step one: Active > 221 DEGs) have a higher number of DEGs (Figure 4C). Results obtained by removing outlier DMSO wells (Step two: Active > 84 DEGs), lowered the minimal number of DEGs to be considered active (Figure 4C).

**Figure 4.**
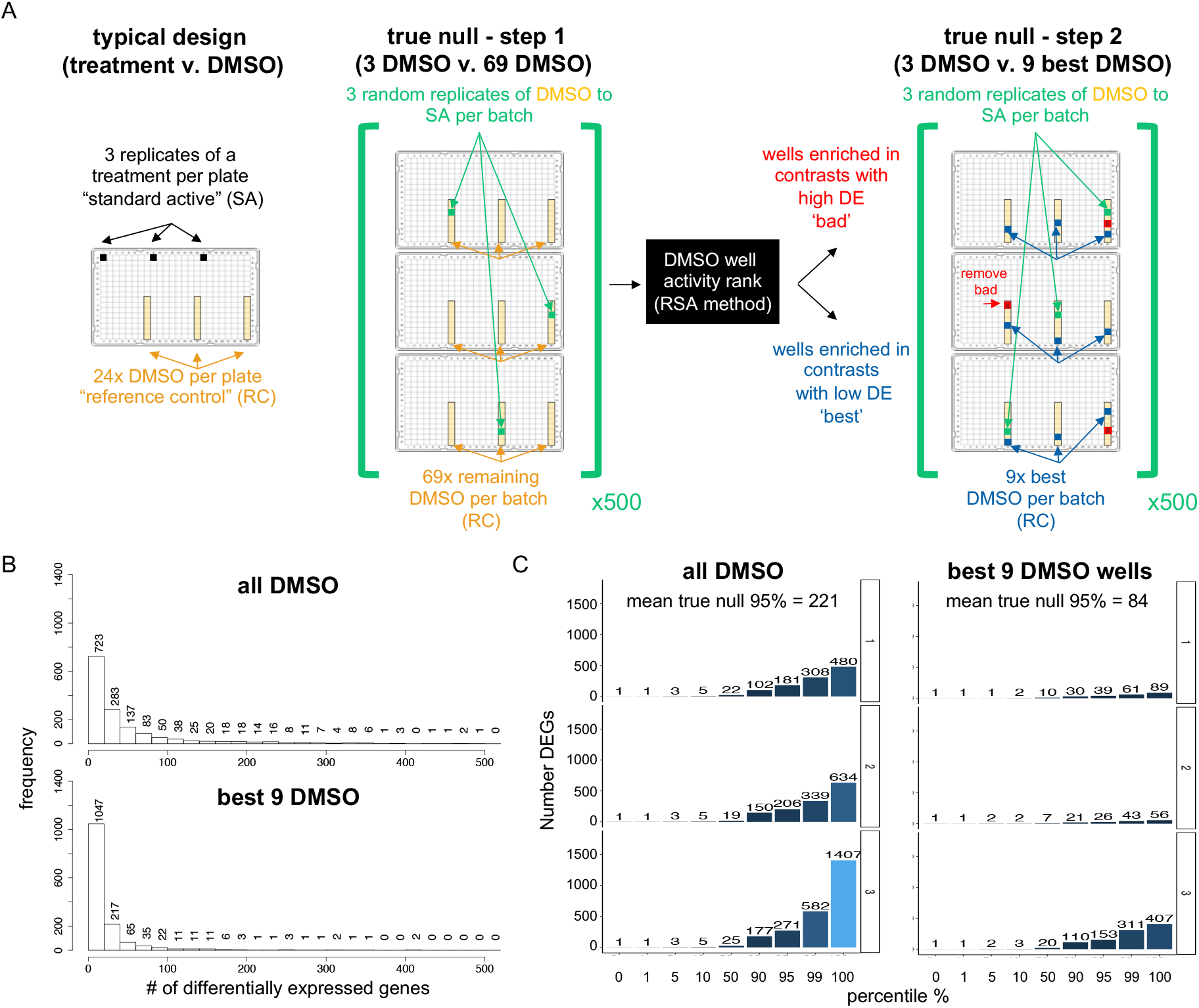
DRUG-seq activity threshold set by the true null calculation. (A) The left panel depicts the typical analysis of a DRUG-seq experiment using compounds with DMSO as a control. When setting a contrast for DE analysis, 3 replicates of a standard active (SA) sample (compound plus dose, colored black) are compared to the DMSO Reference Controls (RC, colored gold)). Each compound treatment has 3 SA well replicates and 24 RC or DMSO well replicates per plate. The middle panel depicts step 1 of the true null calculation. For this, the number of differentially expressed genes is quantified when comparing 3 DMSO well replicates as mock treatments (DMSO/RC turned SA wells, green) relative to the remaining 69 DMSO/RC (gold) wells per batch of 3 plates. 500 randomly chosen differential expression comparisons of 3 DMSO versus the remaining 69 DMSO are performed. Next outlier DMSO wells (colored red) and the best DMSO wells (colored blue) are identified using the Redundant siRNA Activity (RSA) statistical ranking analysis. The right panel depicts step 2 of the true null calculation. 500 random DMSO to DMSO differential expression comparisons are recalculated, this time in the absence of the bad DMSO wells with the 9 best DMSO as the RC. (B) Histogram shows the frequency of the number of DE genes per comparison of randomized DMSO SA to DMSO RC comparisons. In step 1, all DMSO wells are compared (top) and in step 2, the bad DMSO wells are removed and the randomly chosen DMSO wells are compared to the best 9 DMSO wells (bottom). Y-axis is the frequency of the 500 DMSO comparisons, across 3 batches, binned by number of differentially expressed genes on the X-axis. (C) The bar graph plots the number DE genes (y-axis) against the percentile from distribution of 500 randomized DMSO to DMSO comparisons per batch of 3 plates. Left panel depicts true null calculated using all DMSO wells (step 1) and selected a mean threshold of 221 differentially expressed genes (DEG) at a 95 percentile. Depicted on the right panel is the true null calculation removing the outlier DMSO wells across 3 plates (mean 84 DEG at the 95th percentile). The result is interpreted as, only 5% of the time DEGs in DMSO are above the noise detected by comparing DMSO treatments. Removing the outlier DMSO wells for analysis lowers the threshold per batch. Light blue – 1407 DEGs and dark blue is 1 DEG.

For the secondary analysis pipeline discussed below, the step two true null threshold to filter active samples. The secondary analysis pipeline integrates count table data and experimental metadata for quality control and batch correction analyses. In addition to the true null (step two), gene-level thresholds are applied to reduce technical variation. Dimensionality reduction analysis by Uniform Manifold Approximation and Projection (UMAP) (Becht et al., 2019) is then performed to globally visualize the data.

### Batch and Plate Reproducibility Experiment with 14 Tool Compounds

After using the true null threshold to select for active treatments, we used the secondary analysis pipeline and generated a UMAP from the DRUG-seq UMI counts matrix to visualize the global relationship between the 14 compound treatments across an active dose range (Figure 5A). After labelling the UMAP by either Louvain cluster number (Blondel et al., 2008) or compound treatment, it was evident that DRUG-seq identified many transcriptional groups. The 13 Louvain clusters were spatially distinct in the UMAP plot and often displayed substructures in the UMAP projection. We next examined how each compound was distributed across the UMAP and identified that the majority (12 of 14) of compounds were localized in a compound-specific cluster (Figure 5B, File S4, interactive plot). Dilazep and (S)-Crizotinib, were the exception, as these two compounds clustered near DMSO, suggesting low activity in the U2-OS cell line. Some compounds exhibited a single cluster at a certain dose range like BLU9931, CPI-203, CHIR118637 and Cmp_341. For most compounds, dose determined the clustering and we observed dose-specific clusters for Triptolide, Homoharringtonine, Brusatol, NVS-SM2, Fedratinib, Cmp_334 and AZD8055. These compounds may exhibit polypharmacology, and the targets they engage either change and/or interact across doses. Validation would be required to confirm these multi-targeted transcriptional activities. We observed co-clustering of compounds with related MoAs. For example, the translational inhibitors Homoharringtonine and Brusatol exhibited matched dose-dependent clustering, which reflects similar potencies and targets. Cmp_334, a PI3K inhibitor, and AZD8055, an mTOR inhibitor, exhibit a shared cluster at low doses that diverges with increasing concentration. This is not surprising given that PI3K is upstream of mTOR (Hay and Sonenberg, 2004) and perhaps both compounds have selective on-target effects at the lower doses, while a broader range of scaffold-related activity is induced at higher doses. In addition to this descriptive analysis, we quantified cluster composition across batches. We used the Louvain method to define 13 clusters and generated a heat map to indicate the proportion of each active compound treatment within the Louvain clusters per each batch (Figure 5C). Each compound exhibits an enrichment in a specific Louvain cluster and the same result was observed across the 3 batches of cells. This indicates high reproducibility despite both technical and biological variation. Overall, these results indicate that DRUG-seq has sufficient resolution to group compounds by MoA, and because it is target agnostic, it can detect many MoAs in a single assay.

**Figure 5.**
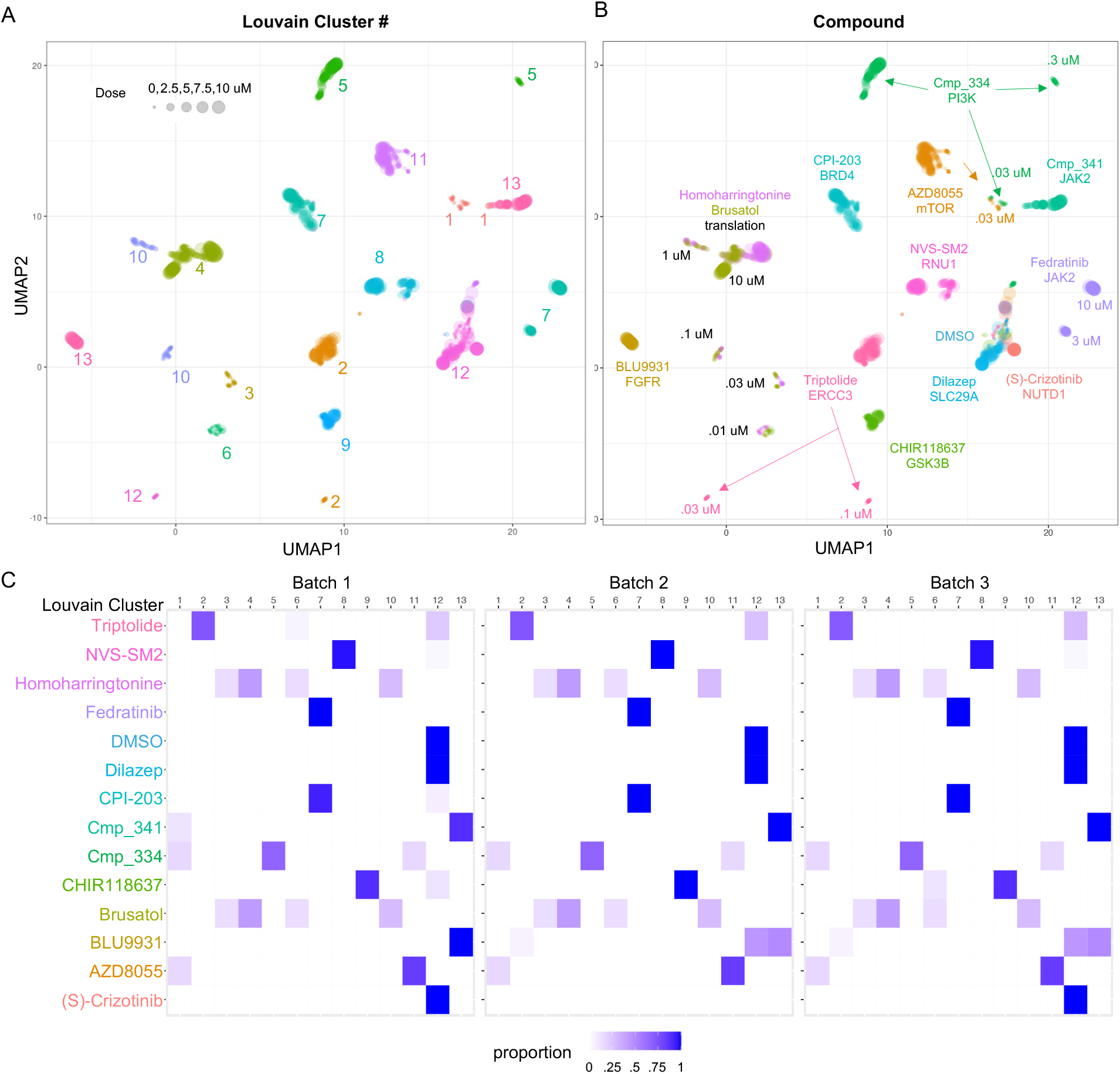
Reproducibility of DRUG-seq data using a set of diverse MoA compounds. (AB) UMAP plots depicting a dimensionality reduced 2D transcriptome using DRUG-seq data colored by Louvain cluster number for A and colored by compound ID for B. The 14-compounds represent a diverse set of MoAs and only active treatments above the true null threshold are plotted and form distinct clusters. Brusatol and Homoharringtonine are both translation inhibitors and co-cluster. DMSO wells are included for an activity reference. Compounds with subtle effects cluster near DMSO. Each dot represents a single well for each treatment. The size of dots is scaled to represent doses from 0-10uM. On the left and right panels each compound or Louvain cluster respectively, is labeled by text in the color of the corresponding dots. (C) Heatmap indicates the proportion of each compound treated sample within Louvain clusters 1-13. Most compounds have a predominant composition in a single or a few Louvain clusters. Cluster 12 includes DMSO and has a mixed composition of many compound and dose combinations of weaker effect. Results across 3 independent batches of cells show a similar composition indicating reproducibility despite biological and technical variation. Proportion for each cluster is indicated with a color scale (0 = white and 1 = dark blue).

Next, the reproducibility of single compounds across doses was examined. Comparing a single dose of Homoharringtonine at 10uM across all 3 batches revealed an average overlap of 68%, which is close to the expected overlap with a false discovery rate threshold of 0.1 (Figure 6A). Homoharringtonine exhibited a dose response and high overlap of DEGs across consecutive doses (Figure 6BC). Within a batch, adjacent doses 1, 3.16, and 10 uM exhibited an average of 82% overlap in differentially expressed genes (Figure 6C). We performed pairwise comparison across all doses and identified a high Pearson’s correlation between samples within a batch. Higher doses and adjacent doses exhibited the highest correlations (.8-.98) (Figure 6D). We generated similar plots for the remaining 13 compounds and observed similar trends in active compounds that clustered away from DMSO. Overall, these studies represent a high bar for vetting the stability of a platform and will allow for comparisons with emerging technologies. We deposited the NGS data, metadata and analysis code for this study to enable other teams to reproduce the transcriptome signatures and to use in additional benchmarking studies.

**Figure 6.**
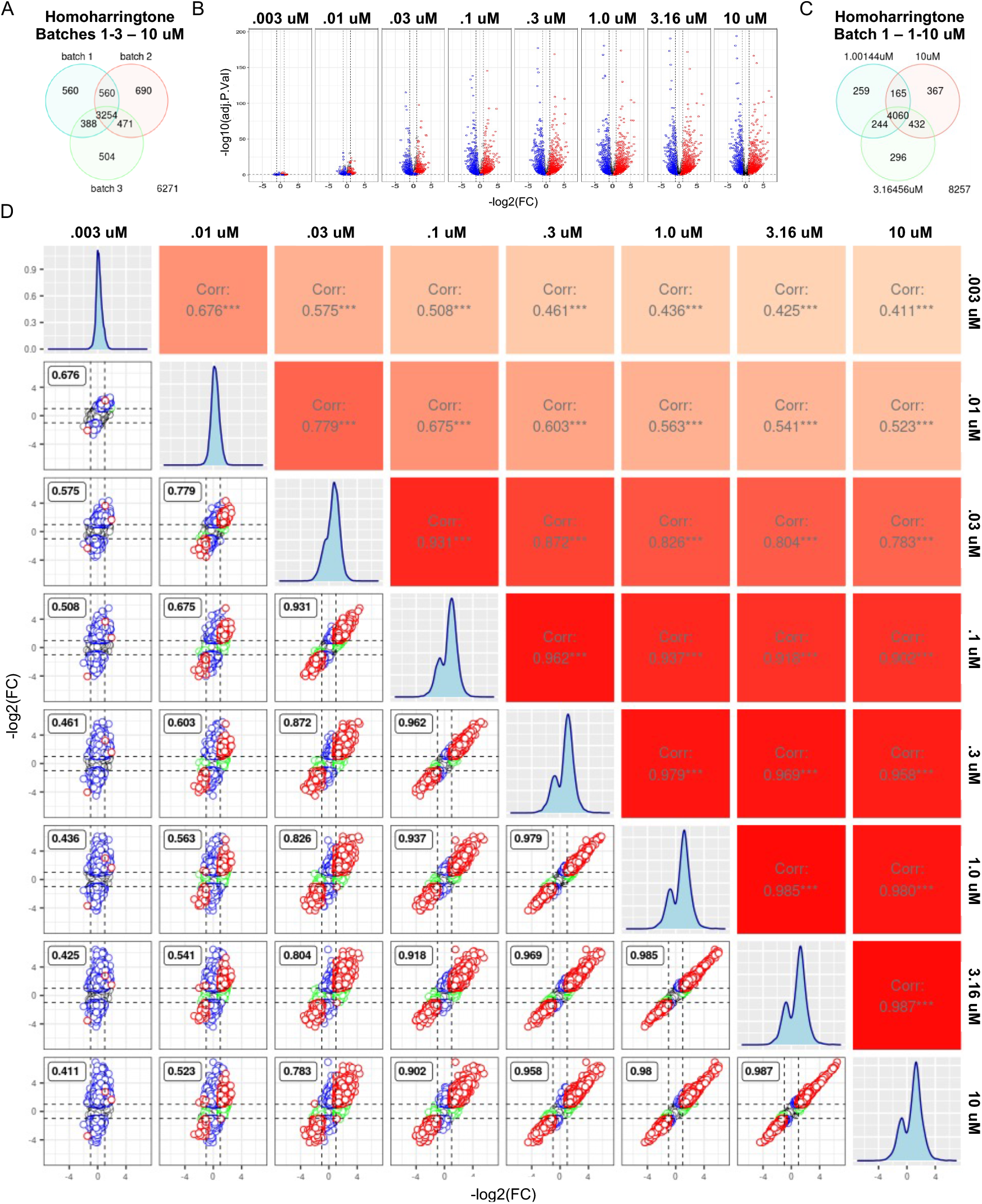
Dose response and batch reproducibility of Homoharringtonine. (A) Venn diagram showing overlapping differentially expressed genes across batches 1-3 at 10uM dose. (B) Volcano plots depicting dose-response for Homoharringtonine. Y-axis adjusted p-value, X-axis log2(FC). Red = upregulated genes 10 uM, Blue = downregulated genes 10uM, n.s. = not significant in 10uM (significant = adj.p<0.5 +/-1 log2FC). (C) Venn diagram depicting overlap of differentially expressed genes between adjacent doses 1-10uM. (D) Pair plot comparing all doses. Top right half indicates Pearson’s correlation between samples, color scale from white = 0 to red = 1. Bottom left scatter plots pairwise compare log2(FC) in gene expression from conditions labeled on the top and right edges. The dashed lines label log2(FC) threshold equal to 1. Red indicates common DEGs across conditions. Blue are DEGs specific to condition on y-axis and green are DEGs specific to the x-axis for each comparison. The histogram on the diagonal the depicts the distribution of gene expression for each condition.

## Discussion

DRUG-seq is a target-agnostic high-throughput screening method with a transcriptome readout, and it can be broadly applied to new cellular models without redesign of the approach or a priori assumptions about key genes or pathways that will be measured. It is well suited for high complexity RNA-seq studies in which many variables and perturbations are tested, such as the dose and length of treatment. DRUG-seq is a bulk RNA-seq readout, and, as such, is best applied to cell cultures with moderate to low heterogeneity. The bandwidth of DRUG-seq accommodates the profiling of a chemical series of related or unrelated chemotypes with different potencies and known or unknown on- and off-target activities. The total cost of DRUG-seq is $3-10 per condition including triplicates at read depth of 0.25 to 1 million reads per well, respectively. This makes it possible to screen thousands of conditions while still providing a high-dimensional readout with greater than 7,000 genes. The resulting high-dimensional data can be used to group compounds by MoA, conduct user-defined signature queries, or search for compounds that may reverse disease signatures. Although the work reported here describes the usage of compounds, one can leverage DRUG-seq profiling for other perturbagens, depending on the question and biological model.

The low cost allowed us to systematically test both the technical and biological variability across plates and batches of cells. By standardizing the experimental design, performance metrics can be tracked long term across many experiments. Using this information, we set statistically defined thresholds to determine the activity of treatments tested in a DRUG-seq experiment. The true null threshold allows us to pick treatments that are statistically defined as active and provides a minimal range of DE genes, which we can trust to produce a reliable signature of expression. We also demonstrated that the results were highly reproducible across plates and batches of cells. The open-source analysis pipeline and available data will facilitate future analytical improvements and lower the barrier for new labs to adopt the platform.

By deploying transcriptomics at scale, we stand to gain biological insights beyond a single target or pathway. With the selection of a set of compounds with diverse MoAs, we demonstrated the granularity of DRUG-seq to discern specific MoAs, dose responses and dose-dependent polypharmacology. This would be easy to miss if only a single or a few doses were tested. DRUG-seq is unbiased, as selection of panels of genes is not required, and it quantifies 5 to 10x more transcripts than L1000 or other targeted amplicon approaches (Li et al., 2012; Subramanian et al., 2017). The wide range of activity detected by DRUG-seq allows it to detect expected and unexpected biological responses. Being blind to the latter has likely contributed to the failure of many drugs at various stages of discovery and development. Furthermore, the dose-dependent detection of on- and off-target phenotypes and the switch between phenotypes can be used to determine potencies for each effect (Renner et al., 2020) and potentially quantify selectivity and safety windows in the absence of dedicated assays.

## Acknowledgements

We would like to thank Dan Palacios and Mark Healy for reviewing the compound lists. We would like to thank Ricardo Dolmetsch for support of the DRUG-seq platform. Thank you, Alan Abrams, for rendering figure 1B.

## Author Contributions

D.J.H., C.Y., and M.H., developed the protocol and generated the 14-compound dataset. A.L. performed sequencing. L.M. tested the protocol as a first-time user and wrote methods. C.R. and M.H. designed the 14-compound experiment. C.R., M.H., C.G.K., K.A.W. and J.L.J. provided experimental feedback. J.L., F.D.S., R.G., M.N., S.R., and D.J.H., developed computational methods to analyze DRUG-seq data. T.T. tested the code as a first-time user. R.J.I., D.J.H., J.L. and K.A.W. wrote the manuscript.

## Additional information

File S1. Metadata for 14-Compound experiment

File S2. Random DMSO wells selected for Step 1

File S3. Random DMSO wells selected for Step 2

File S4. Interactive UMAP (Plotly)

Raw Data: GSE176150

Code: https://github.com/Novartis/DRUG-seq

## Materials and Methods

### Day 1: Cell Culture

U2-OS (ATCC HTB-96) were grown in DMEM, 10% FBS, and 1% Pen/Strep. Sufficient number of cells were grown prior to trypsin dissociation the day of plating. 20 ul of cells were dispensed into 384-well black uClear polystyrene cell culture-treated plates (Griener, Cat#: 781090) using a bottle valve washer/dispenser from the Genomics Institute of the Novartis Research Foundation (GNF, http://www.gnfsystems.com) with a concentration of 5000 cells per well, a day prior to compound treatment. The GNF system is critical for large-scale experiments, but other standard plate wash/dispense equipment or multichannel pipettes will suffice for smaller scale. Density optimization is required for each cell line for optimal downstream steps.

Ngn2 neurons were generated by exposing transgenic H9-hESCs with a dox inducible Ngn2 to doxycycline (1.9 ug/mL) for 3 days in DMEM F12 Glutamax 95%, B27 2%, Pen/Strep 1% and N2 1%. Immature neurons were dissociated using accutase. Ngn2 neurons were then replated in matrigel coated plates at a density of 12,000 per well in 80ul of media (DMEM F12 Glutamax 95%, B27 2%, Pen/Strep 1%, N2 1%, NT3 9.5 ng/mL, 3.8 ng/mL BDNF) with doxycycline (1.9 ug/mL). The Ngn2 neurons were hemi-fed every other day until compound treatment (24h start at day 13).

### Day 1-2: Compound Treatment and Lysis

| **2x Lysis Buffer Stock – store at room temp** |  |
| --- | --- |
| Component | Volume |
| 1M Tris-HCl pH8.0 (Life Tech, 15568-025) | 1000 ul |
| 2M KCl (Ambion, AM9640G) | 750 ul |
| 20% of 20% Ficoll PM-400 in H<sub>2</sub>O (Sigma, F5415-25ML) | 6000 ul |
| Triton-100 (Sigma, T8787-100ML) | 30 ul |
| H<sub>2</sub>O | 2220 ul |

| **2x Lysis Buffer – make fresh each day** |  |
| --- | --- |
| Component | Volume |
| 2x Lysis Buffer Stock | 4800 ul |
| RiboLock RNase Inhibitor 40U/ul (ThermoFisher, EO0381) | 120 ul |

One day after plating, 20 nL of each compound were added using an acoustic dispense Echo 555 Liquid Handler (Labcyte). After a 12 to 24-hour compound treatment, the media was aspirated down to 7.5 ul and an equal amount of 2x lysis buffer was added to all wells using the bottle valve washer/dispenser from GNF (U2-OS Plates were sealed and placed on a microplate shaker HT-91002 (BigBear Automation) for 4 minutes (min) at 900 rpm. Lysis duration is cell type- and density-dependent and requires optimization. Ngn2 neurons plates were lysed for 8 minutes. Plates were then centrifuged at 2000 rpm for 1 min before storage at -80 C until ready for further sample processing. Plates can be stored in –80 C for up to 3-4 weeks, after which RNA quality may begin to deteriorate.

#### Day 3: Reverse Transcription and Library Construction

| **RT Mix** |  |
| --- | --- |
| Component | per well |
| 5XRT Buffer (ThermoFisher, EP0742) | 0.5 ul |
| 1M MgCl<sub>2</sub> (Ambion, AM9530G) | 0.1 ul |
| Regular template switching primer 100 uM (Integrated DNA Technologies) | 0.1 ul |
| RiboLock RNase Inhibitor 40U/ul (ThermoFisher, EO0381) | 0.1 ul |
| dNTP 2.5 uM (ThermoFisher, AM8228G) | 0.1 ul |
| Maxima RT (ThermoFisher, EP0742) | 0.1 ul |
| H<sub>2</sub>O (ThermoFisher, AM8228G) | 1.5 ul |

On the day of sample processing, the assay plates were placed on ice until thawed. In an Armadillo PCR plate (ThermoFisher, AB2384), 2.75 ul of RT mix was added using a Multidrop Combi (ThermoFisher, 5840300). Once the assay plates were thawed, 15 ul of cell lysate was transferred using a Bravo (Agilent Technologies) into the plate containing RT mix. 10 nl of 1 uM barcoded DRUG-seq RT primers were then dispensed into each well using an Echo 555 Liquid Handler (Labcyte). Plates were sealed (BioRad, MSB1001), centrifuged for 1 min at 2000 rpm and incubated at 42 C on a ProFlex PCR system (ThermoFisher, 4484077) for 2 hours.

After RT, samples were centrifuged for 1 min at 2000 rpm. Next, each individual plate was pooled into a reagent reservoir (ThermoFisher, 1064-05-7) using a Bravo Automated Liquid Handling Platform (Agilent). Samples were then transferred from the reservoir into a 50 mL conical tube, purified and concentrated using the DNA clean & concentrator-100 kit (Zymo Research, Cat#: D4030) and eluted in 150 ul of water. Due to the high volume after addition of DNA binding buffer, samples were run three times through the same DNA Clean and Concentrator filter before elution. We further purified the materials eluted from the columns by adding 150 ul (1:1) of AMPure beads RNAclean XP (Beckman coulter, A63987) and incubating for 5 min. The bound beads were pelleted with a magnet and washed twice for 30 seconds (sec) with enough freshly made 80% ethanol to submerge the beads. After removal of ethanol, the beads were allowed to dry completely before eluting with 32 ul of water. To remove single stranded DNA and excess nucleotides, exonuclease I (Exol) treatment was performed on all samples by adding 4 ul ExoI buffer and 4 ul ExoI (New England Biolabs, M0293L). Samples were incubated at 37 C for 30 min, heat inactivated at 85 C for 15 min, and held at 4 C. cDNA was then amplified by adding 50 ul 2X Kapa HIFI PCR ReadyMix (Kapa Biosystems, KK2602), 10 ul of the 10 uM DRUG-seq PCR primer, and then running the following program:

| Temperature | Time | Cycles |
| --- | --- | --- |
| 96 C | Preheat and pause |  |
| 96 C | 1 min | 1 |
| 98 C<br>58 C<br>72 C | 20 sec<br>4 min<br>6 min | 5 |
| 98 C<br>60 C<br>72 C | 20 sec<br>30 sec<br>6 min | 13 |
| 72 C<br>4 C | 10 min<br>Hold (overnight) | 1 |

cDNA samples were purified using the Agencourt RNA clean beads as described above but eluted with 11 ul of water. We ran 1 ul on a Bioanalyzer (Agilent, G2939BA) with DNA high sensitivity chip (Agilent) or Fragment analyzer (Agilent, M5310AA). We expect to see a wide range of fragment sizes, as represented in the figure below. Preamp abundance will be determined by cell type but for U2OS cells, we generally observe quantities of 1 to 5 ng/ul. Size range from 200 to 6000 (representative below).

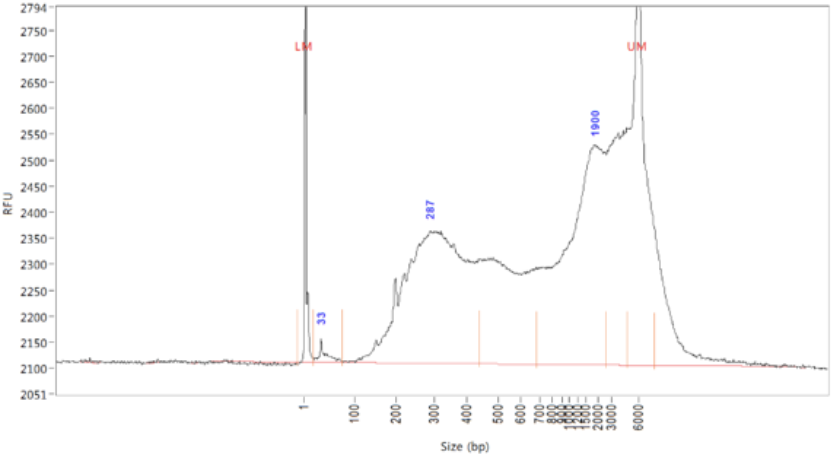

### Day 4: Tagmentation, Purification and Quantification of DRUG-seq libraries

For tagmentation, 5ul of pre-amp material, measured by fragment analyzer, was mixed with nuclease-free water to a final volume of 20 ul. The 20 ul preamp was then mixed with 25 ul TD buffer and 5 ul TDE1 buffer (Nextera kit, FC-131-1096, Illumina) and incubated for 5 min at 55 C and held at 10 C. Tagmented DNA was purified with the Qiagen MinElute PCR Purification Kit (Qiagen, Cat#: 28004) and eluted with 25 ul nuclease-free water.

Each 25 ul sample is then PCR amplified using 15 ul NPM (Nextera XT DNA library preparation kit, FC-131-1024/FC-131-1096), 5 ul DRUG-seq_p5_PCR primer (5 uM) and 5 ul DRUG-seq indexing primer (Table S4). The PCR cycles were as follow:

| Temperature | Time | Cycles |
| --- | --- | --- |
| 72 C | Preheat and pause |  |
| 72 C | 3 min | 1 |
| 95 C | 30 sec | 1 |
| 95 C | 10 sec | 15 |
| 55 C | 30 sec |  |
| 72 C | 30 sec |  |
| 72 C | 5 min | 1 |
| 4 C | Hold |  |

The amplicons were then purified using the Agencourt RNA clean beads as described above but this time eluted with 20ul of nuclease-free water. The samples were then size-selected for 200-600bp fragments using a PippinHT 2% agarose pre-cast gel cassette (Sage Science). 1 ul of the samples from the PippinHT were analyzed on a Fragment Analyzer (Agilent) using a DNF-474 High Sensitivity NGS Fragment Kit 1-6000bp (Agilent, DNF-474-0500). We generally observe approximately 5-10 ng/ul. Averaging a size of 300bp.

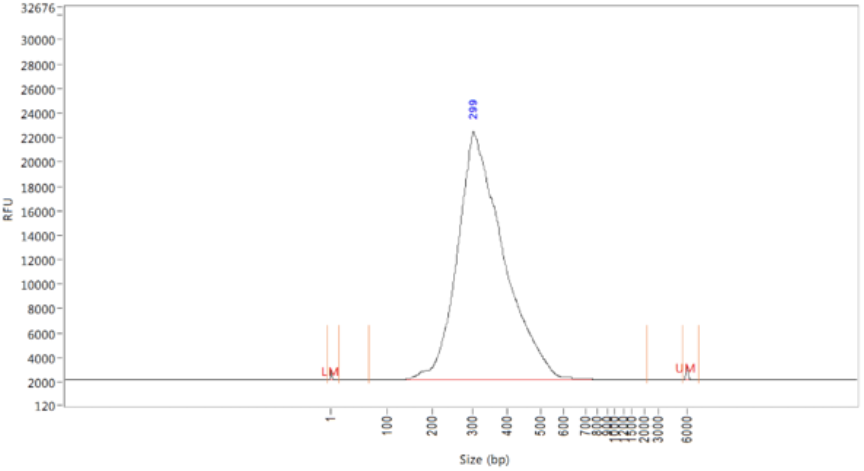

To quantify the libraries, qPCR was performed using Kapa library quantification kit for Illumina (KAPA #KK4824 Roche #07960140001). Following the Kapa kit manual, a pre-mix of 5mL Kapa SYBR FAST qPCR Master Mix, 1mL Illumina primer premix, and 200ul ROX low was combined and the libraries were diluted 1:20,000 with nuclease-free water. The diluted libraries as well as the 6 standards provided in each kit were plated in triplicate using 4ul per well. The reagent pre-mix was plated using 6ul per well. The plate was sealed and run on a QuantStudio 12k Flex with the following cycling:

| Temperature | Time | Cycles |
| --- | --- | --- |
| 95 C | 1 min | 1 |
| 95 C<br>63 C | 15 sec<br>45 sec | 35 |
| 95 C<br>60 C<br>95 C | 15 sec<br>1 min Hold<br>15 sec | Melt Curve Stage |

qPCR data analysis was performed using automatic threshold and baseline settings in the QuantStudio software. The quantities calculated by the software were then dilution corrected by multiplying by the dilution factor above and size corrected following the Kapa kit manual. The libraries were normalized and pooled based on the qPCR quantities. On the following day, library denature was made and sequencing was performed on Illumina’s HiSeq 4000 utilizing a custom Read 1 primer (following manufacturer’s protocol).

### Primer sequences (see Ye et. al., 2018 for full description)

Template switching primer

AAGCAGTGGTATCAACGCAGAGTGAATrGrGrG

DRUG-seq Barcoded RT primers:

AAGCAGTGGTATCAACGCAGAGTACAACAAGGTACNNNNNNNNNNTTTTTTTTTTTT

TTTTTTTTTTTTV

DRUG-seq PCR primer

AAGCAGTGGTATCAACGCAGAGT

DRUG-seq_p5_PCR primer

AATGATACGGCGACCACCGAGATCTACACGCCTGTCCGCGGAAGCAGTGGTATCA

ACGCAGAGT*A*C

DRUG-seq indexing CAAGCAGAAGACGGCATACGAGATNNNNNNNNGTCTCGTGGGCTCGG

DRUG-seq custom read1 primer

GCCTGTCCGCGGAAGCAGTGGTATCAACGCAGAGTAC

## Code and Data

Raw data and processed data: GSE176150 Github: https://github.com/Novartis/DRUG-seq

